# Differentiation and admixture of *Fagus sylvatica* L. and *Fagus orientalis* Lipsky in a northern German forest – learning from pioneer forest work

**DOI:** 10.1101/2022.12.06.519312

**Authors:** Katharina B. Budde, Sophie Hötzel, Markus Müller, Natia Samsonidze, Aristotelis C. Papageorgiou, Oliver Gailing

**Author notes:** Corresponding author: Katharina Budde.

## Abstract

Tree species are suffering from changing and stressful environmental conditions worldwide. *Fagus sylvatica* L., one of the most common Central European deciduous tree species showed symptoms of crown damage, a reduction in growth and increased mortality following the severe recent drought years. For Germany *Fagus orientalis* Lipsky, a closely related species with higher drought tolerance, originating from south-eastern Europe, Turkey, the Greater Caucasus region and the Hyrcanian forest, has been proposed as an alternative with high future potential. The translocation of pre-adapted planting material has been proposed as a tool to mitigate negative effects of climate change. This approach can be beneficial but might also harbor risks.

Taking advantage of *F. orientalis* trees planted over 100 years ago in the forest district of Memsen, Germany, we set out to study admixture between the two beech species and the direction of gene flow. Furthermore, we used a range-wide dataset of *F. sylvatica* and *F. orientalis* provenances to determine the origin of the introduced trees. Using a combination of nuclear EST-SSRs and one chloroplast SSR marker with species-specific variants, we could show that interspecific gene flow was going in both directions. In most cases, *F. sylvatica* was the pollen donor which is likely explained by the higher abundance of this species producing vast amounts of pollen. The planted trees originated from the Greater Caucasus region and showed strong genetic divergence from German *F. sylvatica* populations. In the future, gene flow patterns as well as hybrid performance from different provenances should be tested in additional stands and in comparison to *F. sylvatica* provenances from southern Europe to assess the suitability of Oriental beech for the mitigation of climate change impacts.

## Introduction

Currently, environmental conditions are changing worldwide and the consequences are visible in many tree stands. If natural stands are not locally adapted anymore due to a change in e.g. temperature and precipitation regimes and they are not able to respond at the individual physiological level (plastic response), they have to adapt (shift in allele frequencies over generations), migrate (gene flow via seed and pollen dispersal) to places with more suitable conditions or go extinct (Aitken et al., 2008). In the last years, we have already seen an increased mortality due to more prolonged or extreme summer droughts in tree species in Central Europe (Schuldt et al., 2020). This shows that individuals less adapted to the new conditions are currently being selected out. Typically, tree stands show high phenotypic plasticity, a high standing genetic variation upon which selection can act, and many tree species exhibit efficient gene flow (Petit and Hampe, 2006). However, if tree populations are able to adjust to the ongoing fast change in environmental conditions by gene flow and shifting allele frequencies is still unclear (Alberto et al., 2013; Chevin et al., 2010; Sork, 2016). Additionally, cultural landscapes in most parts of the terrestrial world present a rather restricted matrix for migration to places with more suitable conditions, in particular for species with specialized resource and habitat requirements (Tscharntke et al., 2012). And in tree species, their long generation times in which evolution is happening on time scales that are not tangible for humans produces uncertainty about the future of our forest ecosystems.

In forestry, for the use of planting material, alternative species or provenances of the same species, naturally growing in other regions with similar environmental conditions to the conditions expected in the future under climate change in the target region, are being discussed (Pötzelsberger et al., 2020). For Germany, recently, nine tree species were recommended with high future potential that should be tested in provenance trials for their suitability for silviculture in Germany and under future conditions (Liesebach et al., 2021). The tree species listed by the expert working group included among others also exotic species, such as Caucasian fir (*Abies nordmanniana* (Stev.) Spach), Oriental beech (*Fagus orientalis*), Atlas cedar (*Cedrus atlantica* G. Manetti) and Turkish hazel (*Corylus colurna* L.). For some other exotic species, which are planted in Germany and have potentially a high suitability for planting under future environmental conditions, such as Douglas fir (*Pseudotsuga menziesii* (Mirb.) Franco) or Japanese larch (*Larix kaempferi* (Lamb.) Carr.), there is already much experience present regarding provenance choice and hence less intensive testing is foreseen (Liesebach et al., 2021). The introduction of exotic species for forestry purposes has a long tradition worldwide. In central Europe, several economically important tree species were introduced in the last centuries, e.g. *P. menziesii* (Knoerzer and Reif, 2002) and *Quercus rubra* L. (Nicolescu et al., 2019) from North America. Most of the introduced species, do not pose a threat to the natural local ecosystems, however some became invasive, such as *Robinia pseudoacacia* L. which can now overgrow and form monospecific stands in sites with previously species-rich dry and semi-dry grasslands (Vítková et al., 2017). Compared to species introductions, assisted gene flow (AGF, translocation of potentially pre-adapted planting material of autochthonous species but from other regions of the distribution range) is supposed to harbor fewer risks. One potential effect of newly introduced taxa or lineages is hybridization and introgression into closely related local populations. Gene flow and introgression between allochthonous and local populations can be beneficial, as it might introduce new alleles with positive effects (Aitken and Whitlock, 2013) and has been recommended for species with low adaptive potential in the face of global warming (Aitken and Bemmels, 2015). However, it can also pose risks, such as outbreeding depression and the disruption of local adaptation to non-climatic factors and changes in the population composition and structure (Ellstrand et al., 1999; Laikre et al., 2010). The effects of exotic gene flow on local populations are of particular importance for populations of conservation concern such as small, relict populations. This has been studied, e.g., in *Populus nigra* L. introgressed by widely planted North American *P. deltoides* W. Bartram ex Marshall (Cagelli and Lefevre, 1995; Rathmacher et al., 2010; Vanden Broeck et al., 2004), *Pinus pinaster* Aiton (Unger et al., 2016, 2014) and Eucalypt species in vicinity of plantations of exotic Eucalypt species (Barbour et al., 2010, 2008). However, a systematical assessment of conservation concerns prior to the introduction of new provenances or tree species is lacking.

Oriental beech, one of the nine mentioned high potential tree species for future silviculture in Germany originates from south-eastern Europe and extends to the Caucasus and the Hyrcanian forest in Iran. If Oriental beech and European beech (*F. sylvatica*) should be considered subspecies or distinct species, is still under debate (Gömöry et al., 2018; Renner et al., 2016). They show strong differentiation and diverged ca. 8.7 (20–1.8) Ma ago (Cardoni et al., 2022; Renner et al., 2016). Both taxa show differences in habitat preference, with *F. orientalis* growing typically on warmer and drier sites. Here, we took advantage of a beech stand in Lower Saxony, northern Germany, in which the local forester, Friedrich Erdmann, introduced *F. orientalis* trees over 100 years ago. The planted trees are now growing intermixed in a *F. sylvatica* stand with abundant natural regeneration. Both species or subspecies are known to hybridize in their sympatric distribution range, e.g. in Greece (Müller et al., 2019; Papageorgiou et al., 2008). We were interested if the two taxa were able to cross and form viable offspring in Germany. To this end, we collected leaf samples from adult and young trees of the two taxa in the forest district of Memsen and used nuclear EST-(expressed sequence tags) and chloroplast microsatellites to assess the interspecific gene flow. We specifically evaluated if gene flow was occurring in both directions and if we could identify backcrosses. Additionally, we assessed the usefulness of morphological leaf traits to differentiate the two taxa in Germany and determined the origin of the planted *F. orientalis* trees.

## Material and methods

### Plant material

Leaves from a total of 115 *F. sylvatica* and *F. orientalis* trees were collected in two plots (ca. 14 km apart) in the forest district Memsen located ca. 50 km south from the city of Bremen in Germany in summer 2021. Around 100 years ago individuals of *F. orientalis* were planted that are nowadays as big and vital as the *F. sylvatica* trees growing next to them. We collected leaves from 23 adult *F. orientalis* trees (DBH > 10 cm) that we could find, as well as from some of the offspring morphologically resembling *F. orientalis*. Additionally, we collected 22 reference samples from *F. sylvatica* adult trees and also offspring growing in direct vicinity. As reference, we gathered previously published genotypic data of nine microsatellites from nine *F. sylvatica* populations collected in Germany and Greece and 13 *F. orientalis* stands from Turkey and Iran (Bijarpasi et al., 2020; Müller et al., 2019). Furthermore, we added one more newly collected, stand from the Republic of Georgia. The stand is located in the Lagodekhi nature reserve at the southern slopes of the Caucasus, in the region of Kakheti (2000-2500 meters above sea level). Leaves were gathered from a total of 50 trees at a minimum distance of ca. 30 m. Our final data set comprised 698 trees from 24 locations (Fig. 1, Table S1, Supplementary Material).

**Figure 1.**
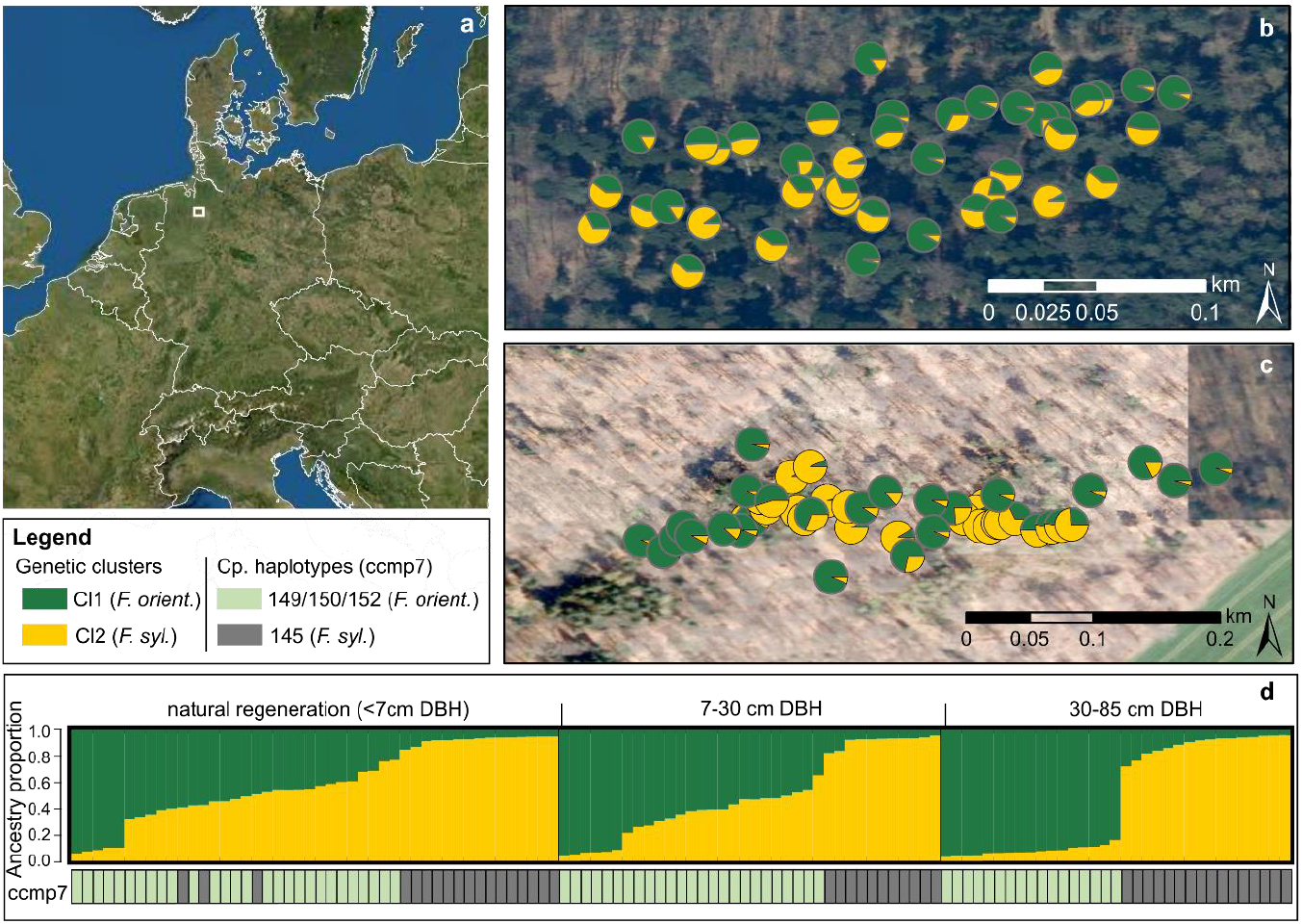
**(a)** Overview map of Germany indicating the location of the forest district Memsen. **(b)** and **(c)** Pie charts showing the individual ancestry proportions of two genetic clusters for each beech tree in the two study sites in Memsen. **(d)** Barplot of ancestry proportions for two genetic clusters ordered by height class and below a color bar indicating the chloroplast haplotype of *ccmp7*. As indicated in the legend, dark green depicts the genetic cluster representing *F. orientalis* and yellow the one of *F. sylvatica*. Chloroplast haplotypes 149, 150 and 152 (light green) are typical for *F. orientalis*, while 145 (gray) originates from *F. sylvatica*.

### Morphological data

For a total of 83 trees (all trees sampled on the second of a total of two collection days), three leaves were collected, dried and fixed on a sheet of paper. We scanned the leaves and used the software *Easy leaf area* (Easlon and Bloom, 2014) to determine the leaf area. Additionally, the following leaf morphological traits were assessed: number of leaf veins, leaf lamina length, petiole length, the lamina width measured from the central vein to the leaf margin at the widest point of the lamina. We estimated the leaf shape ratio as LSR=100xWP/LL and the leaf-petiole ratio as PR=100xPL/(LL+PL).

### DNA extraction and genotyping

Leaves for DNA extraction (115 leaves from Memsen and 50 leaves from Georgia) were collected in paper bags and dried using silica gel orange (Roth, Karlsruhe, Germany). Approximately 1cm^2^ (ca. 10-20 mg) of dried material per leaf was ground to powder using a Retsch mill (Retsch, Haan, Germany). The DNeasy 96 Plant Kit (Qiagen, Hilden, Germany) was used following the manufacturer’s instructions for DNA extraction. All DNA stocks were 1:10 diluted prior to PCR amplifications. All samples were genotyped at nine EST-SSR markers (FgSI0006, FgSI0009, FgSI0024, FS_C1968, FS_C2361, FS_C4971, FS_C6785, FS_C7377, FS_C7797) published previously (Burger et al., 2018; Kubisiak et al., 2009) following the procedure described in Müller et al. (2019). Additionally, one chloroplast microsatellite, *ccmp7* (Weising and Gardner, 1999) that was known to differentiate the two taxa in Greece and Turkey (Gailing and von Wuehlisch, 2004; Hatziskakis et al., 2009) was also amplified using the same protocol. Fragment sizes were determined on an ABI 3130xl Genetic Analyzer (Applied Biosystems, Foster City, USA) against GS 500 ROX (Applied Biosystems, Foster City, USA) as internal size standard. Allele scoring was conducted with the GeneMapper 4.0 software (Applied Biosystems, Foster City, CA, USA). Some samples previously genotyped in Müller et al. (2019) were also included in Bijarpasi et al. (2020) and additionally in the new PCR and genotyping runs to ensure that all data sets yielded comparable genotypes and could be analyzed jointly.

### Data analyses

#### Species differentiation and introgression in a mixed stand in Memsen, Germany

First, we checked our multilocus genotypes from the forest district in Memsen for genotype duplicates resulting from root suckers and one sample had to be removed. Afterwards we used the admixture model with correlated allele frequencies implemented in the Bayesian clustering algorithm in STRUCTURE v. 2.3.4 (Pritchard et al., 2000) to identify *F. sylvatica, F. orientalis* and admixed individuals. We tested K=1 to K=6, with 10 independent runs per K, a burn-in length of 100,000 and a run length of 200,000 simulations. The results were uploaded to the STRUCTURE harvester (Earl and von Holdt, 2012) and CLUMPAK (Kopelman et al., 2015) pipelines for the evaluation of summary results. As expected, the Evanno approach based on ΔK (Evanno et al., 2005) in combination with the mean log likelihood of K, as well as a visual inspection of the barplots indicated that K=2 was well explaining our data. We compared the cluster assignment of reference samples (adult trees, clearly identified as *F. sylvatica* and *F. orientalis* based on morphological traits) and the *ccmp7* chloroplast haplotype (145 bp typical for *F. sylvatica* and 150 bp and 152 bp indicative for *F. orientalis*) with the cluster assignment to identify which cluster was representing which species. One adult tree showed the chloroplast haplotype 149 bp at *ccmp7* and this individual could clearly be assigned to *F. orientalis* based on nuclear SSRs.

Comparing the STRUCTURE cluster assignment proportions with the DBH of *F. orientalis* trees, showed that several individuals with an assignment proportion between 0.84 and 0.875 for the genetic cluster corresponding to *F. orientalis* also had a DBH of >40 cm. As it is unlikely that a third generation backcross (typically expected assignment proportion of ca. 0.875) already exhibits such a big DBH, we assumed that these trees must rather belong to the originally planted *F. orientalis* trees and defined 0.84 as the threshold for clear assignment to one of the two clusters. All trees showing intermediate assignment proportions (<0.84 and >0.16) were considered hybrids. Afterwards, the multilocus genotype data was read in R v. 4.1.2 (R Developmental Core Team, 2021) and converted to a genind object using *adegenet* (Jombart and Ahmed, 2011). We performed a Principal Components Analysis (PCA) using the function *dudi.pca* while coloring the individuals based on their STRUCTURE cluster assignments as described above to confirm the genetic structure in Memsen.

#### *Morphological differentiation of* F. sylvatica *and* F. orientalis *in Memsen*

The leaf morphological data was read in R v. 4.1.2 (R Developmental Core Team, 2021). First, we tested for correlations between leaf morphological traits using Spearman’s correlation coefficients (Table S2, Supplementary Material). We removed highly correlated traits showing correlation coefficients above 0.7 and retained four traits (leaf area, number of leaf veins, petiole length and lamina-shape ratio) for further analyses. These four traits were used to perform a PCA. The first two principal components were plotted and visually compared with the genetic species assignment by coloring the individuals based on the genetic cluster assignments. Additionally, boxplots were created for each morphological trait to highlight the differences between *F. sylvatica*, *F. orientalis* and their hybrids.

#### *Range wide genetic structure and the origin of the* F. orientalis *trees planted in Memsen*

We selected the adult trees from Memsen that were clearly identified as *F. sylvatica* and *F. orientalis* and combined their genotypic data with the genotypes from Lagodekhi (Georgia) and previously published data from Müller et al. (2019) and Bijarpasi et al. (2020). This yielded a total data set of 698 individuals (Table S1) from 25 populations, genotyped at nine microsatellite markers. First, we checked the number of individual multilocus genotypes and removed 20 duplicated genotypes. Genetic diversity parameters, such as the number of private alleles (*A*_P_), mean rarified allelic richness over loci based on 16 allele copies (*A*_R_), expected heterozygosity (*H*_e_) and the inbreeding coefficient (*F*_IS_) with 95% confidence intervals based on 1000 bootstraps were estimated for each population using the *hierfstat* package (Goudet, 2005) in R. Additionally, we computed pair-wise population differentiation as *G*_ST_ (Nei, 1973; Nei and Chesser, 1983) implemented in the R package *mmod* (Winter, 2012).

Then, we ran STRUCTURE using the same model settings as described above but assessing clusters from K=1 up to K= 16 for the range-wide dataset. Again, the CLUMPAK and STRUCTURE harvester pipelines were used to evaluate the STRUCTURE results. Furthermore, we performed a PCA as described above using the range-wide genotypic data in R.

Pairwise genetic differentiation (*G*_ST_) between the four genetic clusters identified by STRUCTURE was assessed by assigning all populations to one of the four clusters (Table 1). Only Hilia from Greece was excluded from this analysis as this stand showed considerable admixture between clusters.

**Table 1.** Genetic diversity estimates per sample location for 24 *Fagus sylvatica* (FS) and *F. orientalis* (FO) populations and the *F. orientalis* trees with unknown origin from Memsen. ID, population number; gCluster, genetic cluster assignment based on STRUCTURE results; N, number of samples; *A*_P_, number of private alleles; *A*_R_, mean rarified allelic richness over loci based on 16 allele copies; SE, standard error; *H*_o_, observed heterozygosity, *H*_e_; expected heterozygosity and *F*_IS_, inbreeding coefficient; 95% CI, 95% confidence interval after bootstrapping.

| ID | Population | Country | Species | gCluster | N | $A_P$ | $A_R$ (SE) | $H_e$ (SE) | Fis [95% CI] |
| --- | --- | --- | --- | --- | --- | --- | --- | --- | --- |
| 1 | Memsen | Germany | FS | CI1 | 31 | 8 | 3.098 (0.474) | 0.446 (0.092) | -0.016 [-0.095, 0.129] |
| 2 | Goehrde | Germany | FS | CI1 | 24 | 0 | 2.941 (0.388) | 0.410 (0.085) | -0.032 [-0.089, 0.032] |
| 3 | Calvoerde | Germany | FS | CI1 | 24 | 0 | 3.078 (0.384) | 0.474 (0.060) | -0.042 [-0.148, 0.040] |
| 4 | Aetomilitsa | Greece | FS | CI1 | 24 | 1 | 3.541 (0.593) | 0.539 (0.060) | -0.030 [-0.101, 0.049] |
| 5 | Tsepelovo | Greece | FS | CI1 | 24 | 0 | 3.366 (0.468) | 0.530 (0.068) | 0.034 [-0.125, 0.161] |
| 6 | Alevitsa | Greece | FS | CI1 | 24 | 0 | 3.317 (0.565) | 0.510 (0.065) | -0.007 [-0.071, 0.080] |
| 7 | Varnuntas | Greece | FS | CI1 | 24 | 0 | 3.428 (0.482) | 0.523 (0.067) | -0.021 [-0.112, 0.088] |
| 8 | Frakto | Greece | FS | CI1 | 24 | 0 | 3.346 (0.434) | 0.514 (0.057) | -0.030 [-0.143, 0.089] |
| 9 | Lepida | Greece | FS | CI1 | 24 | 0 | 3.356 (0.426) | 0.560 (0.046) | **0.115 [0.040, 0.217]** |
| 10 | Hilia | Greece | FO/FS | CI1/CI2 | 24 | 0 | 3.759 (0.537) | 0.595 (0.054) | 0.043 [-0.098, 0.195] |
| 11 | Demirkoy | Turkey | FO | CI2 | 24 | 1 | 3.697 (0.611) | 0.510 (0.072) | **-0.127 [-0.206, -0.018]** |
| 12 | Catalca | Turkey | FO | CI2 | 10 | 0 | 3.683 (0.779) | 0.500 (0.091) | 0.062 [-0.107, 0.299] |
| 13 | Inegoel | Turkey | FO | CI2 | 10 | 1 | 4.154 (0.776) | 0.546 (0.091) | **0.139 [0.005, 0.272]** |
| 14 | Izmit | Turkey | FO | CI2 | 10 | 2 | 3.927 (0.597) | 0.515 (0.088) | -0.053 [-0.097, -0.008] |
| 15 | Duezce | Turkey | FO | CI2 | 10 | 0 | 3.774 (0.531) | 0.526 (0.080) | -0.06 [-0.229, 0.125] |
| 16 | Covakici | Turkey | FO | CI2 | 10 | 0 | 3.081 (0.296) | 0.446 (0.075) | -0.024 [-0.108, 0.085] |
| 17 | Karabuek | Turkey | FO | CI2 | 10 | 0 | 3.758 (0.464) | 0.548 (0.063) | -0.065 [-0.197, 0.046] |
| 18 | Lagodekhi | Georgia | FS | CI3 | 49 | 13 | 3.167 (0.640) | 0.397 (0.078) | 0.072 [-0.027, 0.182] |
| 19 | Bolor_W | Iran | FO | CI4 | 47 | 1 | 3.319 (0.766) | 0.395 (0.098) | -0.012 [-0.085, 0.125] |
| 20 | Bolor_E | Iran | FO | CI4 | 49 | 2 | 3.200 (0.640) | 0.394 (0.095) | 0.037 [-0.025, 0.114] |
| 21 | Siyahroud_W | Iran | FO | CI4 | 47 | 0 | 2.785 (0.530) | 0.337 (0.093) | **0.087 [0.021, 0.142]** |
| 22 | Siyahroud_E | Iran | FO | CI4 | 45 | 2 | 3.243 (0.586) | 0.388 (0.084) | 0.037 [-0.062, 0.077] |
| 23 | Changool_W | Iran | FO | CI4 | 43 | 5 | 3.221 (0.670) | 0.364 (0.096) | -0.029 [-0.055, -0.004] |
| 24 | Changool_E | Iran | FO | CI4 | 43 | 1 | 3.119 (0.590) | 0.394 (0.087) | -0.048 [-0.115, 0.009] |
| 25 | Unknown (Memsen) |  | FO | CI3 | 23 | 28 | 3.165 (0.676) | 0.379 (0.098) | -0.019 [-0.161, 0.123] |

## Results

### Species differentiation and admixture in Memsen

The analyses of the population genetic structure of beech trees in Memsen identified two distinct genetic clusters, representing *F. sylvatica* and *F. orientalis*, respectively (Figure 1). Also admixed individuals were revealed in both sample locations in Memsen (Figure 1b and c) and chloroplast marker *ccmp7* showed that both species, *F. sylvatica* and *F. orientalis*, could serve as mother trees in interspecific crosses (Figure 1d). Most commonly the *F. orientalis* chloroplast alleles were found in admixed individuals indicating that in most of these cases *F. orientalis* was the mother tree. We differentiated three size classes. The biggest trees (DBH 30-85 cm) could all be assigned to one of the two species, *F. sylvatica* and *F. orientalis*, respectively. However, in the intermediate and small size classes many individuals with varying levels of admixture were identified. The PCA based on the nuclear microsatellite genotypic data from Memsen confirmed the genetic structure (Figure S1). Genetic differentiation between trees clearly assigned to *F. orientalis* and *F. sylvatica* in Memsen was strong (*G*_ST_ =0.292, *P*< 0.001).

The four leaf morphological traits showed clear differences between the two beech species growing in Memsen and hybrids exhibited intermediate phenotypes (Table S2 and Figure S2, Supplementary Material). *Fagus orientalis* was characterized by bigger leaves, with more leaf veins and shorter petioles compared to *F. sylvatica*. A PCA based on the four uncorrelated leaf traits coincided well with the genetic species assignment (Figure 2).

**Figure 2.**
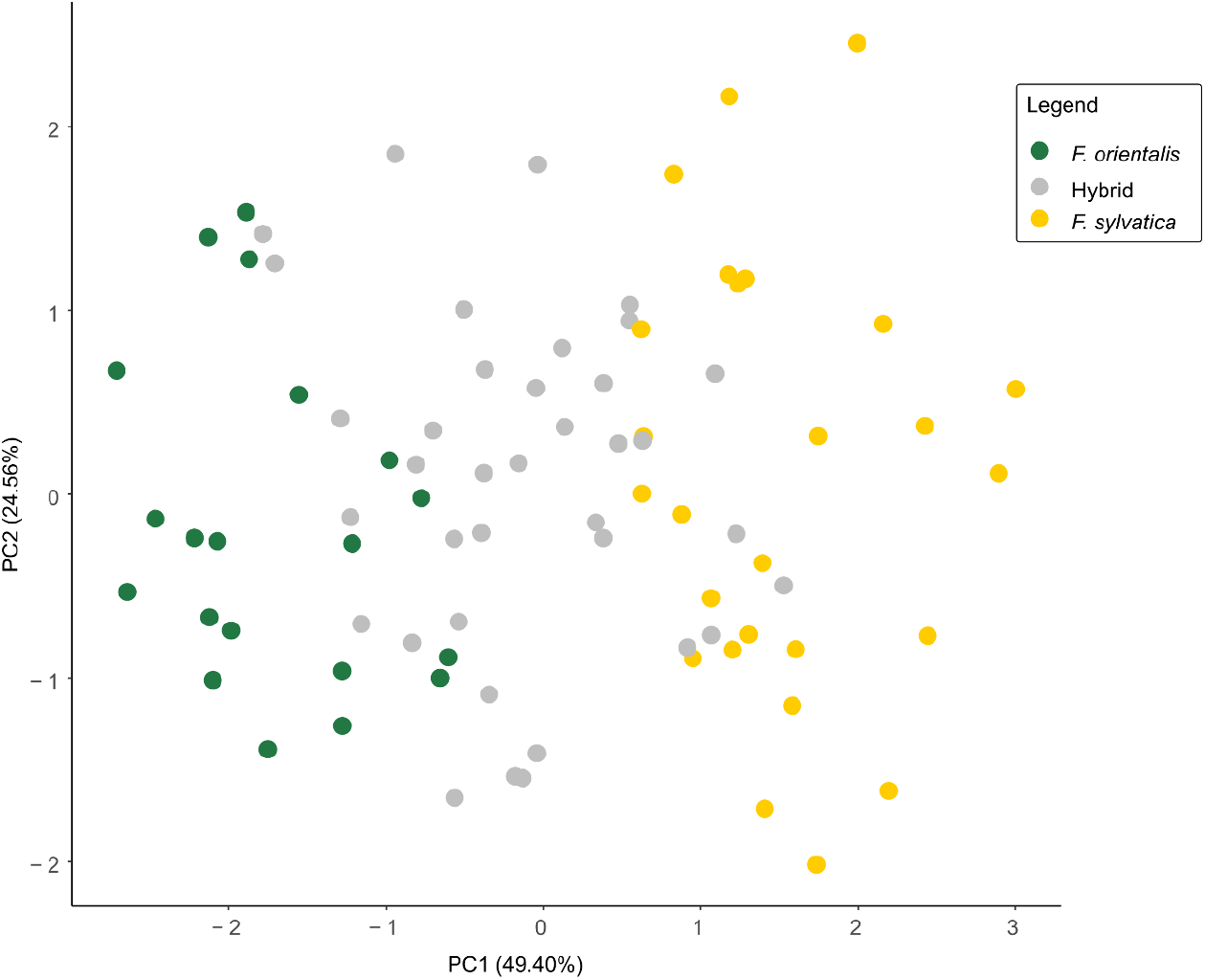
Principal components analyses of leaf morphological data *of F. orientalis, F. sylvatica* and admixed samples from Memsen. The coloration is based on the genetic assignment from the STRUCTURE analyses with admixed samples shown in gray.

### *Origin of the* F. orientalis *trees planted in Memsen and range-wide population genetic structure*

The range-wide genotypic data revealed four gene pools. *Fagus sylvatica* trees from Germany and Greece represented the first genetic cluster (Cl1), the second to fourth genetic clusters comprise *F. orientalis* trees originating from Turkey (Cl2), Georgia (Cl3) and Iran (Cl4), respectively (Figure 3). One population, Hilia from Greece, indicated admixture between the two species (between Cl1 and Cl2) occurring here in sympatry in the natural range. The *F. orientalis* trees planted in Memsen could clearly be assigned to the third gene pool (Cl3) represented by samples from Georgia.

**Figure 3.**
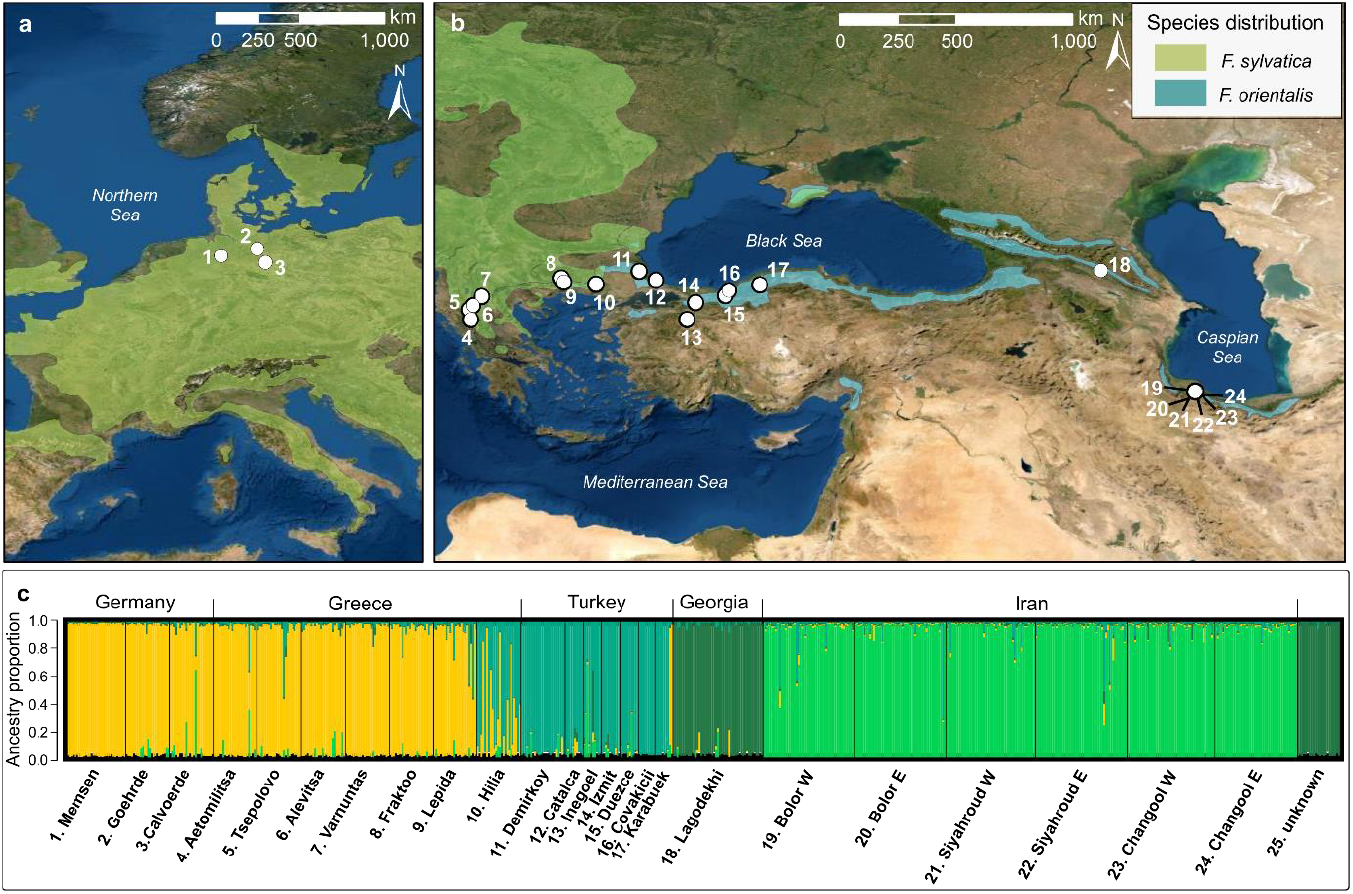
**(a)** and **b)** Overview maps showing the origin of *Fagus sylvatica* and *Fagus orientalis* samples. **(c)** Barplot of ancestry proportions of four genetic clusters for samples from range-wide beech populations. The *F. orientalis* trees growing in the forest district Memsen are labeled as “25. unknown” and are not depicted in the maps.

The genetic diversity, e.g. allelic richness and expected heterozygosity were highest in the admixture zone of the two species in Greece and Turkey and slightly lower in Germany (*F. sylvatica*), Iran and Georgia (*F. orientalis*). Lagodekhi from Georgia and the *F. orientalis* trees from Memsen showed the highest numbers of private alleles. Differentiation between the four gene pools, assessed as pairwise *G*_ST_ was high ranging from 0.130 to 0.200 (Table 2) and was of course lower between sample locations within gene pools (Table S3).

**Table 2.** Genetic differentiation, assessed as pair-wise *G*_ST_ estimates, between the four genetic clusters of *F. sylvatica* and *F. orientalis*. All values are significant with *P*<0.001. Cl1, *F. sylvatica* populations from Germany and Greece (excluding Hilia); Cl2, *F. orientalis* populations from Turkey; Cl3, *F. orientalis* populations from the Caucasus region; Cl4, *F. orientalis* populations from Iran.

|  | CI1 | CI2 | CI3 | CI4 |
| --- | --- | --- | --- | --- |
| CI1 |  | 0.130 | 0.200 | 0.139 |
| CI2 |  |  | 0.186 | 0.106 |
| CI3 |  |  |  | 0.188 |

## Discussion

In the current situation where forest practitioners are considering alternative species for reforestation efforts after certain tree species are suffering under the changing climatic conditions, we would like to call for caution. In this study, we could show that *F. orientalis* trees planted over 100 years ago in Northern Germany were growing well in the new habitat and hybridized with autochthonous *F. sylvatica* trees growing in direct vicinity. *Fagus orientalis* has been recommended as species with high future potential in Germany but there is an urgent need for prior research initiatives confirming the suitability of *F. orientalis* provenances and a careful assessment of the risks.

Hybridization between *F. orientalis* and *F. sylvatica* has previously been reported in the natural range where both species co-occur (Müller et al., 2019; Papageorgiou et al., 2008) and also in locations where *F. orientalis* was planted in Central Europe (Kurz et al., 2022). Using a combination of EST-SSRs and the chloroplast SSR *ccmp7*, we could show that gene flow is going in both directions. We observed F1 hybrids but also backcrosses which had to be expected in this old (>100 years) beech stand assuming a generation time of 40-50 years (Comps et al., 2001). Only three introgressed saplings derived from *F. sylvatica* as seed parent, while in all other hybrids *F. sylvatica* was the pollen donor. However, a quantitative assessment of the direction of gene flow between the two species in Memsen was hampered by the unequal abundance of the two species. *Fagus sylvatica* is the main stand forming species, producing vast amounts of pollen while only 18 big *F. orientalis* trees (with DBH > 20cm) were present in Memsen. Flowering phenology in both species is metandric with female flowers of one tree being receptive before pollen release on the same tree and controlled pollination experiments between both species previously reported good success rates (Nielsen and Schaffalitzky de Muckadell, 1954). Kurz et al. (2022) observed that spring phenology (assessed on buds and leaves) was slightly earlier in *F. orientalis* compared to *F. sylvatica* in two sites in Switzerland suggesting that this might also be the case for the flowering phenology. A similar finding in spring phenology was observed in a growth chamber experiment, comparing populations of the two species from their natural range in Greece (Varsamis et al., 2019). Based on this we could potentially expect a better overlap between pollen releases of *F. orientalis* with female flower receptivity of *F. sylvatica* than *vice versa*. In any case, the abundance of hybrids in Memsen and other central European stands (Kurz et al., 2022) shows that there is a substantial overlap in flowering phenology enabling interspecific gene flow.

The four genetic clusters detected in range-wide *F. sylvatica* and *F. orientalis* samples coincide with the genetic structure detected in previous studies (Bijarpasi et al., 2020; Cardoni et al., 2022; Kurz et al., 2022; Müller et al., 2019). The *F. orientalis* trees planted in Memsen were assigned to Cl3 and therefore most likely originate from the Greater Caucasus region. Only the *F. orientalis* trees from Memsen and the population from Lagodekhi represent Cl3. Differentiation between the two populations was low but significant (*G*_ST_=0.053; *P*<0.05) and both populations harbored the highest numbers of private alleles. This most likely reflects that the gene pool Cl3 was least well represented in our samples but also that the *F. orientalis* trees from Memsen do not originate from the direct vicinity of Lagodekhi. Kurz et al. (2022) also identified the Greater Caucasus region as the main source of planting material for *F. orientalis* in Central Europe. Genetic differentiation between the two species in Memsen was high, which was also reflected by the leaf morphological traits enabling species determination.

The translocation of pre-adapted planting material to improve climate adaptation within a species range (Aitken and Whitlock 2013) is a necessary tool for conservation, particularly for certain populations under existential threat, i.e. where levels of genetic diversity are so low that they are likely to be unable to adapt to changing conditions (Ennos et al., 1998; Swarts and Dixon, 2009). However, especially, when introducing genetically strongly divergent provenances, AGF can also pose risks, such as outbreeding depression and might disrupt local adaptation. *Fagus sylvatica* is one of the most abundant deciduous tree species in Central Europe and a foundation species in many European forest ecosystems (Leuschner and Ellenberg, 2017). It harbors high levels of genetic variation (e.g. Vornam et al. 2004; Buiteveld et al. 2007; Piotti et al. 2012; Rajendra et al. 2014) and although damaged *F. sylvatica* trees have been reported after the recent drought years (Schuldt et al., 2020), typically not all trees within a stand showed the same level of damage and many remained healthy (Pfenninger et al., 2021). Recent studies therefore suggest that there is considerable variation at adaptive traits, including drought resistance within stands (Cuervo-Alarcon et al., 2021; Pfenninger et al., 2021; Pluess and Weber, 2012). Furthermore, natural regeneration after drought events is a key factor and is abundant in *F. sylvatica*. Early life history traits in *F. sylvatica* are highly plastic and emphasize the ability of the species to acclimatize quickly at the short term (Muffler et al., 2021; Müller et al., 2020). In a provenance test, where seedlings from several Greek *F. sylvatica* and *F. orientalis* populations were tested in a growth chamber under simulated conditions reflecting climate change scenarios, dramatic plastic responses that allowed the survival of seedlings were also recorded in bud burst, leaf senescence and growth rate of both species (Varsamis et al., 2019). Additionally, Beloiu et al. (2020) could show that *F. sylvatica* saplings defoliated during the drought in 2018 exhibited a high recovery rate in 2019. However, mortality under dry and hot conditions is high (Muffler et al., 2021) and in the long term, especially with increasing temperatures and longer drought periods, *F. sylvatica* populations will need to genetically adapt ideally facilitated by the high genetic variation within stands.

## Conclusions

Here, we could show that *F. sylvatica* and *F. orientalis* hybridized in an old growth mixed beech forest in Northern Germany and both species could act as pollen and seed parent. This interspecific gene flow can be beneficial, harboring potential for AGF. However, the need, usefulness and risks of the introduction of *F. orientalis* planting material with the aim of AGF should carefully be evaluated. We second the proposal of Kurz et al. (2022) to take advantage of already introduced *F. orientalis* plantations in Central Europe to assess the impact of gene flow from different *F. orientalis* provenances on neighboring *F. sylvatica* stands. This could shed light, e.g. on the risk of outbreeding depression on adaptive traits and the performance of admixed seedlings and saplings could be assessed under controlled environmental conditions of potential future environmental scenarios. Furthermore, if AGF will be deemed necessary for *F. sylvatica* we propose that also less divergent *F. sylvatica* provenances from drier regions such as e.g. Spain, Rumania or Greece could be suitable candidates for AGF and should also be evaluated. Bert et al. (2020) studied the suitability of different *Quercus petraea* (Matt.) Liebl. and *Q. robur* L. provenances for translocation, by assessing their performance in three common garden sites with contrasting environmental conditions in northern France. They could show that the suitability of provenances needs to i be carefully evaluated and non-climatic factors such as e.g. soil type also need to be taken into account.

## Acknowledgements

We acknowledge the help of Mr. Uwe Niedergesäss from the forest district Memsen in Hoyershagen who granted permission for sampling and Dr. Lars Drößler who facilitated sample collection in Lagodekhi. Furthermore, we would like to thank Dr. Victor Chano for his help during sample collection and Alexandra Dolynska for help in the laboratory.

## Data availability

All data will be made available upon acceptance on Zenodo

## Author contributions

Katharina B Budde: Conceptualization, Investigation, Data Curation, Formal analysis, Supervision, Writing – original Draft Sophie Hötzel: Investigation, Data Curation, Formal analysis Markus Müller: Data Curation, Writing – Review & Editing Natia Samsonidze: Investigation, Writing – Review & Editing Aristotelis C. Papageorgiou: Writing – Review & Editing Oliver Gailing: Conceptualization, Supervision, Writing – Review & Editing

## Supplementary Material

**Table S1** Overview of range wide *Fagus sylvatica* and *F. orientalis* sample locations, GPS coordinates and *N*, the number of samples collected in each location.

**Figure S1** Plot of the first two principal components derived from the genotypic data of nine microsatellites for all *F. sylvatica* and *F. orientalis* samples collected in Memsen, Germany. The coloration is based on the genetic assignment from the STRUCTURE analyses with admixed samples shown in gray.

**Table S2** Pearson correlation coefficients of leaf morphological traits of *F. sylvatica* and *F. orientalis* trees collected in Memsen. Significance levels: *, *P*<0.05; **, *P*<0.01; ***, *P*<0.001; n.s.: not significant. nu., number; r., ratio.

**Figure S2** Boxplots of four morphological traits from leaves of *F. sylvatica, F. orientalis* and admixed individuals growing in Memsen, Germany. The coloration is based on the genetic assignment from the STRUCTURE analyses with admixed samples shown in gray.

**Figure S3** Plot of the first two principal components based on the genotypic data of nine microsatellites for all range wide *F. sylvatica* and *F. orientalis* samples. The coloration depicts the country where the samples were collected.

**Table S3** Pair-wise genetic differentiation (*F*_ST_) between *F. sylvatica* and *F. orientalis* sample locations. unkn., unknown

(provided as separate file).

