## Supplementary material for "Differentiation and admixture of *Fagus sylvatica* L. and *Fagus orientalis* Lipsky in a northern German forest – learning from pioneer forest work"

**Table S1** Overview of range wide *Fagus sylvatica* and *F. orientalis* sample locations, GPS coordinates and *N*, the number of samples collected in each location.

| Sample location | Country | *N* | Species | Latitude | Longitude |
|---|---|---|---|---|---|
| Goehrde | Germany | 24 | *F. sylvatica* | 53.123 | 10.820 |
| Calvoerde | Germany | 24 | *F. sylvatica* | 52.404 | 11.261 |
| Memsen | Germany | 32 | *F. sylvatica* | 52.767 | 8.8990 |
| Aetomilitsa | Greece | 24 | *F. sylvatica* | 40.273 | 20.813 |
| Tsepelovo | Greece | 24 | *F. sylvatica* | 39.883 | 20.876 |
| Alevitsa | Greece | 24 | *F. sylvatica* | 40.432 | 20.963 |
| Varnuntas | Greece | 24 | *F. sylvatica* | 40.807 | 21.328 |
| Frakto | Greece | 24 | *F. sylvatica* | 41.545 | 24.524 |
| Lepida | Greece | 24 | *F. sylvatica* | 41.386 | 24.622 |
| Hilia | Greece | 24 | *F. orientalis* | 41.302 | 25.934 |
| Demirkoy | Turkey | 24 | *F. orientalis* | 41.820 | 27.662 |
| Catalca | Turkey | 10 | *F. orientalis* | 41.467 | 28.350 |
| Inegoel | Turkey | 10 | *F. orientalis* | 39.883 | 29.600 |
| Izmit | Turkey | 10 | *F. orientalis* | 40.567 | 29.950 |
| Duezce | Turkey | 10 | *F. orientalis* | 40.850 | 31.150 |
| Covakici | Turkey | 10 | *F. orientalis* | 41.050 | 31.283 |
| Karabuek | Turkey | 10 | *F. orientalis* | 41.283 | 32.533 |
| Bolor-W | Iran | 50 | *F. orientalis* | 37.001 | 50.073 |
| Lagodekhi | Georgia | 50 | *F. orientalis* | 41.830 | 46.330 |
| Bolor-E | Iran | 50 | *F. orientalis* | 36.996 | 50.073 |
| Siyahroud-W | Iran | 49 | *F. orientalis* | 36.993 | 50.094 |
| Siyahroud-E | Iran | 50 | *F. orientalis* | 36.990 | 50.090 |
| Changool-W | Iran | 48 | *F. orientalis* | 36.981 | 50.069 |
| Changool-E | Iran | 46 | *F. orientalis* | 36.977 | 50.073 |
| Memsen | unknown origin | 23 | *F. orientalis* | - | - |

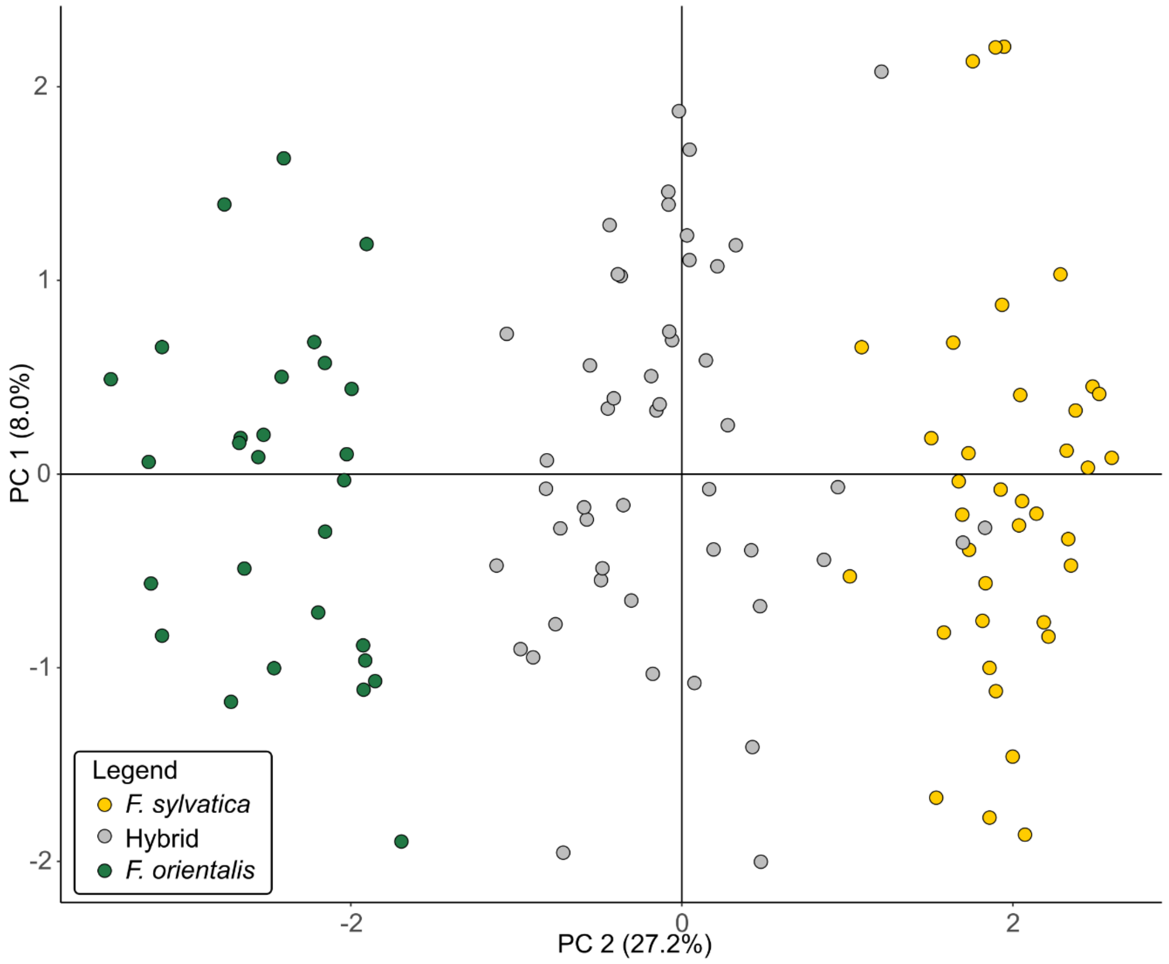


**Figure S1** Plot of the first two principal components derived from the genotypic data of nine microsatellites for all *F. sylvatica* and *F. orientalis* samples collected in Memsen, Germany. The coloration is based on the genetic assignment from the STRUCTURE analyses (q-values) with admixed samples shown in gray.

**Table S2** Pearson correlation coefficients of leaf morphological traits of *F. sylvatica* and *F. orientalis* trees collected in Memsen. Significance levels: *, *P*<0.05; **, *P*<0.01; ***, *P*<0.001; n.s.: not significant. nu., number; r., ratio.

| | leaf area | n. of leaf veins | leaf length | petiole length | lamina width | Lamina-shape r. | leaf-petiole r. |
|---|---|---|---|---|---|---|---|
| leaf area | | 0.57*** | 0.89*** | -0.14n.s. | 0.84*** | -0.24* | -0.47*** |
| nu. of leaf veins | | | 0.66*** | -0.30** | 0.50*** | -0.44*** | -0.56*** |
| lamina length | | | | -0.12n.s. | 0.81*** | -0.46*** | -0.51*** |
| petiole length | | | | | 0.08n.s. | 0.36** | 0.90*** |
| lamina width | | | | | | 0.13n.s. | -0.26* |
| lamina-shape r. | | | | | | | 0.51*** |
| leaf-petiole r. | | | | | | | |

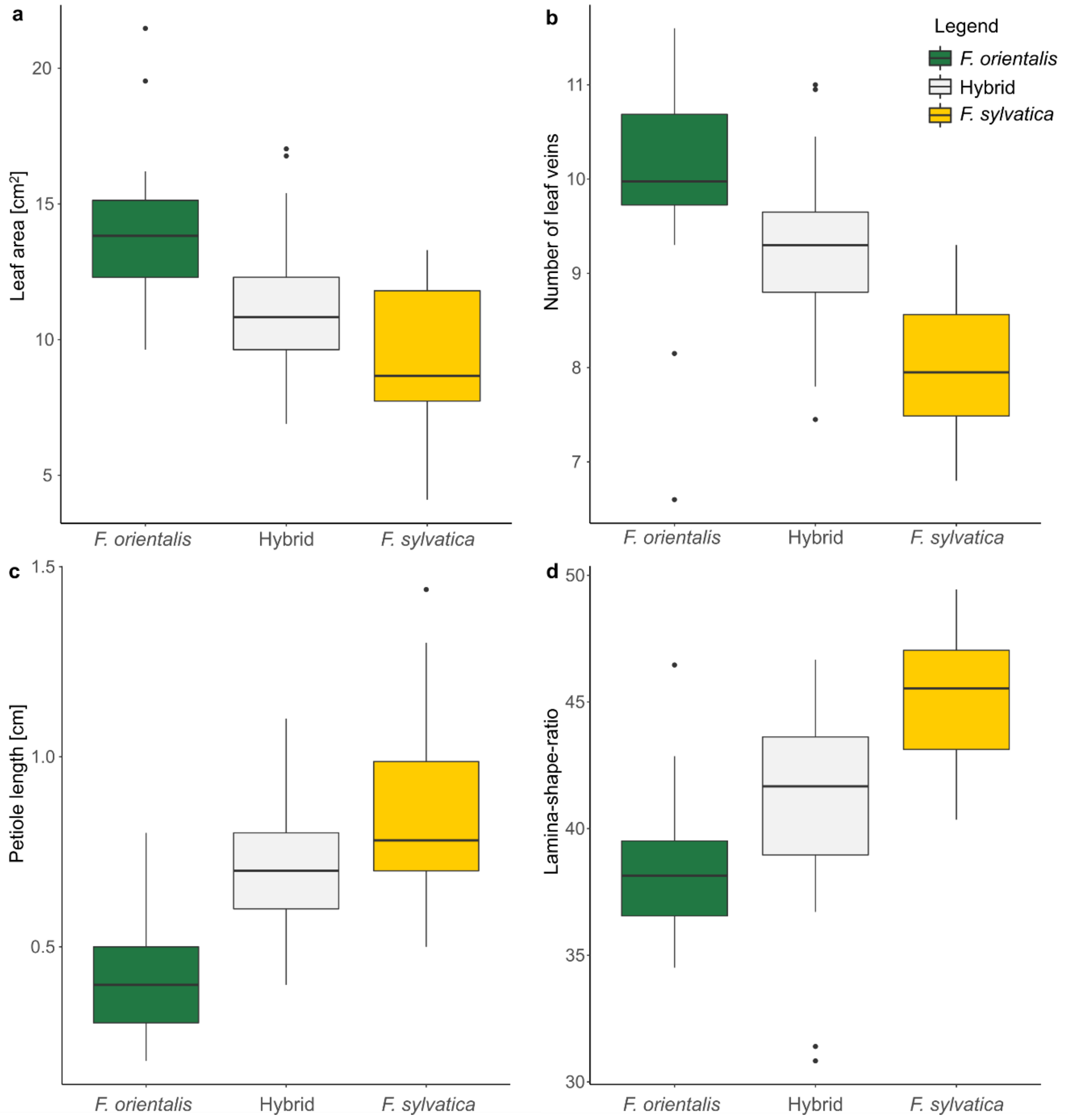


**Figure S2** Boxplots of four morphological traits from leaves of *F. sylvatica*, *F. orientalis* and admixed individuals growing in Memsen, Germany. The coloration is based on the genetic assignment from the STRUCTURE analyses with admixed samples shown in gray.

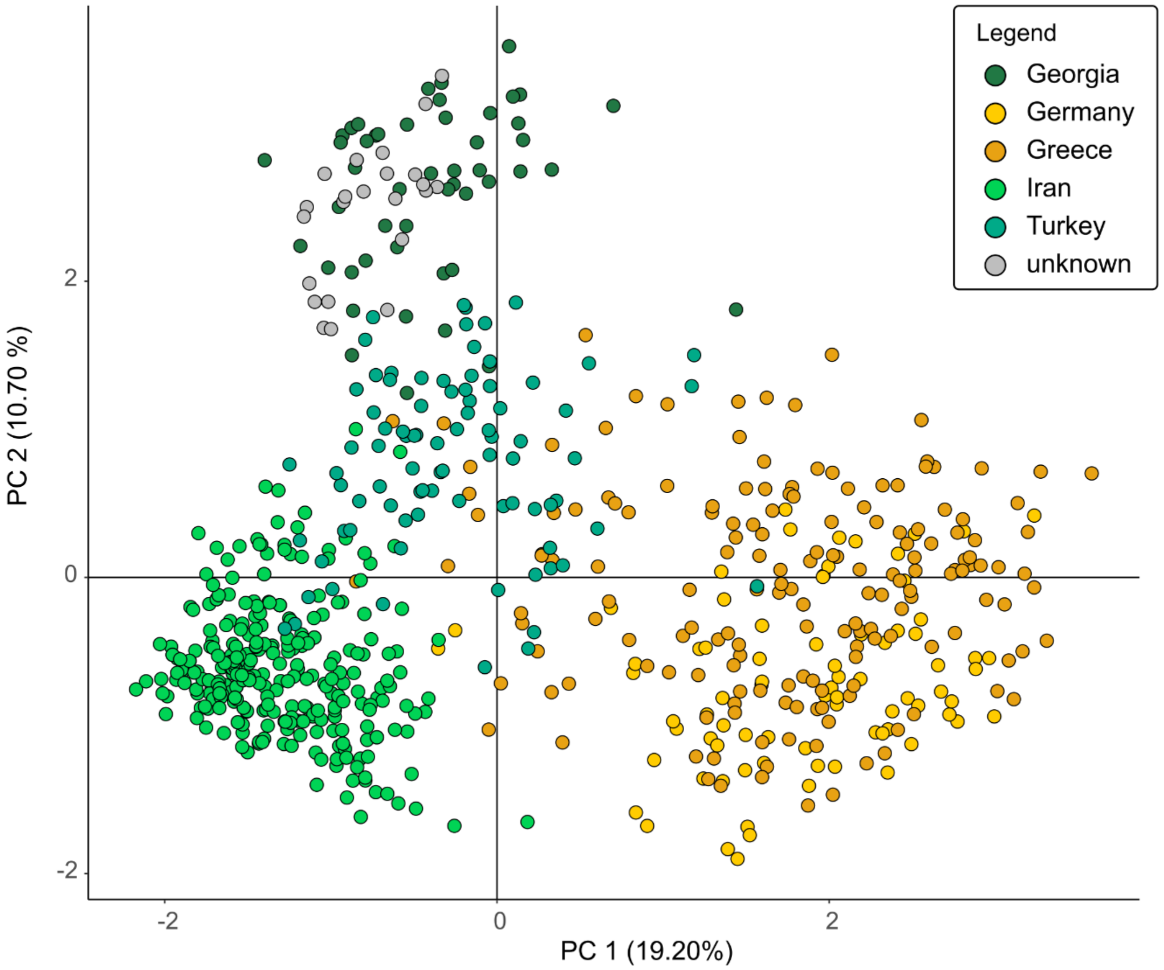


**Figure S3** Plot of the first two principal components based on the genotypic data of nine microsatellites for all range wide *F. sylvatica* and *F. orientalis* samples. The coloration depicts the country where the samples were collected. The planted *F. orientalis* trees from Memsen are shown in gray (unknown).

**Table S3** Pair-wise genetic differentiation ($F_{ST}$) between *F. sylvatica* and *F. orientalis* sample locations. unkn., unknown
(provided as separate file).
